# Therapy Induced Tumor Senescence Model

**DOI:** 10.1101/2022.10.23.513380

**Authors:** Ghanendra Singh

**Affiliations:** iCurious.in

## Abstract

Senescent cell accumulation and defective clearance of the senescent cells by the immune system occur with aging and increase the prevalence of diseases like cancer. Anti-tumor therapies can induce senescence in the tumor cells. Senescence Associated Secretory Phenotypes (SASP) secretion by these senescent tumor cells activates the innate NK cells which can detect and eliminate them. Mechanisms are unclear about how does it occur? A combination of immunotherapy and senotherapy has shown the possibility to reduce the tumor burden and increase the health span. The temporal and intensity dynamics of the therapeutic dose regimen remains to be studied. Therefore, a simplified therapy-induced senescence (TIS) phenomenological model is proposed to explain the mechanism of senescent tumor cell clearance by the NK immune cells and understand the possibility of a two-punch therapy technique in regulating tumors. Interaction strength changes for the cellular population within a healthy and an aged tumor microenvironment. The simulation result shows an oscillatory behavior existing between the tumor and immune cells. Tumor heterogeneity acts as inherent noise in sustaining the tumor for relapse emergence despite therapeutic clearance. The model indicates the formation of a robust oscillatory loop between the tumor, immune, and senescence cells which they can tune by modifying the phenotypic fitness landscape through secreted factors making them resistant despite selective removal of the sensitive populations by various therapies. The model highlights the importance of modified and aged tumor microenvironment by senescence tumor cells in obstructing clearance of both senescence and tumor cells by the innate immune system. Cancer therapies along with senolytics may have a robust and effective regulatory potential over tumor and senescence cells. The model also provides a preliminary analysis of the therapy temporal and intensity dosage regimen causing a therapeutic shift in tumors.

## Introduction

Cellular senescence refers to a cell cycle arrest state due to replication stress, ROS, DNA damage, telomere shortening or other cellular stress. It was identified as a replication limit in the somatic cells after certain number of divisions [1]. Different type of senescnce induction [2] exists namely: replicative senescence [1], stress induced senescence [3], oncogene-induced senescence [4], and lastly therapy-induced senescence (TIS) [5]. Cancer cells in therapy induced senescence (TIS) are stress tolerant and can survive for longer duration. Cancer cells in TIS modify their microenvironment which favors senescence maintenance, escape, and proliferation of senescence cells. Apart from tumor cells modulating immune cells, senescent cells can also attract and modulate the immune cells that eliminate them. Senescence cell accumulation and immune dysfunction are related with each other.

Senescent cell accumulation threshold theory state that once senescence cell burden exceeds a threshold, the self-amplifying paracrine and endocrine spread of senescence through SASP outpaces the clearance of senescence cells by the immune system [6], it may impede its function [7] and can accelerate other aging mechanisms [8]. Senolytic interventions can delay its onset and prevent age-related diseases and the onset of cancer [9]. Potential therapies could delay, prevent, or alleviate age-related disorders by targeting mechanisms linking senescent cells to immunological dysfunction [7]. Cellular senescence is also induced by anticancer therapies and they undergo a switch from antitumor to protumor role via paracrine signaling and cell-intrinsic mechanisms. Hence, effective modulation of the senescence associated secreted phenotype (SASP) can improve the efficacy of therapies [10].

Cancer cells can escape from senescence [11]. Cancer cells initially respond to the therapy and can persist as dormant tumor cells relapsing at a later stage. Senescence in cancer cells due to therapy possibly a form of dormancy [12]. Cancer cells undergoing therapy induced senescence (TIS) may be at the origin of tumor relapse. An increasing body of evidence from preclinical studies demonstrates that radiation and chemotherapy cause the accumulation of senescent cells both in tumor and normal tissue. Senescence cells in tumors can paradoxically promote tumor relapse, metastasis, and resistance to therapy in part through expression of the SASP.

Therapy induced senescence (TIS) is a strong phenotypic adaptation which modifies the fitness and may provide a control strategy for robust therapeutic population level control for tumor as in the case of evolutionary double bind approach [13]. Two punch therapy technique [14] have similar parallels to the double bind.

Predatory effects of the immune system over sensitive tumors such as in immunotherapy and activation of the natural killer cells for elimination of the senescent cancer cells along with senolytics theoretically if possible may hold the key to selectively control and eliminate tumor burdon and clearing senescence cancer cells. It may prevent phenotypic adaptation in tumor for longer periods. Indicating predator facilitaiton [13] from applied ecology to utilize multiple predator to control resistant tumor population. TIS may place the senescence cancer cells in evolutionary double bind by utilizing two punch therapy.

Combining inhibitor drugs induces senescence phenotype in tumor naturally activating natural killer (NK) cells for immune control system [15]. SASP helps in recruiting and activating immune cells to clear senescence cells [16]. Its important to understand the mechanisms of senescence tumor cell elimination by immune cells. Is cytokines are involved? Does it protects non senescence tumor cells? [17]. Senolytics can be used to target cancer cells in TIS. It reduced cancer relapse in mice treated with doxorubicin, suppress SASP, and reduce anti-cancer treatment dosage [18]. Possibility of introducing senolytic in combination with radiotherapy, immunotherapy such as two punch therapy [14].

Adaptive therapy [19] approaches have shown good results in regulating tumor population through competitive advantage and utilizing phenotypic cost. Another proposed therapeutic approach is targeting senescent cells using one-two punch cancer therapy. Cancer therapies (first punch) induce senescence both in tumor and normal tissue. Selective clearance using senotherapeutics (second punch) in tumors will prevent relapse, metastasis, and resistance to therapy. Dosage timing of second punch therapy is extremely important to improve the efficacy [14]. Reflects the importance of understanding therapy schedule, and dosage timing to achieve effective results in the regulation of tumor and senescence cell burden.

Classical predator prey formulation of cancer immune system was given by [20]. Model consists of three equations for tumor, effector, and IL-2 cytokines [21]. Lotka Volterra model can also be used to understand the interaction between cells within tumor microenvironment [22]. It can explains the growth, death, and interaction between populations. Another model of tumor sub population dynamics during therapy [23]. Dynamics of the system can explain the immunobiology states such as elimination, equilibrium, and escape states of the cancer [24]. Tumor immuno-surveillance hypothesis formulated in 1957 indicates that the immune system is capable of inhibiting the growth of very small tumors and eliminating them before they become clinically evident. It relates well with the threshold theory of senescence cells [6].

A tumor’s immunogenic phenotype is constantly shaped by the tumor microenvironment. Several models have been proposed to explain tumor immune dynamics. A mechanistic mathemamtical model which describes the interactions between tumor and immune system was proposed by [25]. Following are also three dimensional models [26, 27]. Another model explained the tumor and immune cell dynamics [28]. Discussed the role of mathematical modeling of radiotherapy along with tumor-immune interactions. Formation of positive inflammmatory feedback loop between senescence and immune system in aging niche [29]. Phenotypic characteristic of senescence cells with initiation through cell cycle arrest, oncogene activation, or DNA damage. Early senescence with progressive chromatin remodeling, and late senescence with inflammaging [30]. Existence of two states in non immune senescence cells. Immunogenic cells secrete SASP factors promoting senescence clearance. Anergic cells do not stimulate the immune responses. Due to aging or external stimuli such are secreted from tumor leading to switching from anergic state to immunogenic state. Another model for chemo-immunotherapy based treatment strategies [31].

In this article, proposed model consider therapy induced senescence (TIS) in tumor at the base. It does not consider the competitive advantage of sensitive population over resistance sub population and phenotypic heterogeneity seen in tumor populations just to keep it simple to test the hypothesis offered by the therapy induced senescence (TIS) landscape. Taking inspiration from TIS and simplify the system to consider three cellular populations namely: tumor cells (T), immune cells (I), and senescence cells (S) within aged tumor microenvironment may provide some interesting insights. Initially start with a deterministic model involving three interacting entities and later model its stochastic dynamics. First (i), model focus on the importance of eliminating the dormant tumor cells [32]. Second (ii), it explains the mechanism of senescence tumor cell elimination by NK cells. Third (iii), it try to explain the proposed senescence cell accumulation threshold hypothesis [6]. Fourth (iv) role of senescence cells in modifying the TME pre and post therapy. Fifth (iv), therapy dosage intensity and temporal dynamics for TIS with combined immunotherapy and senotherapy. Lastly, investigate the aspects of tumor, immune, and senescence cells interaction during therapy and discuss the importance of testing the hypothesis for experimental validation in modeling the aged tumor immune microenvironment.

## Methods

Python for writing the ODEs, plotting nullclines and bifurcation plots. Earlier classic model [20] and the review by [25] is used as a baseline to construct the model.

## Results

### Tumor Immune Senescence Model

Modeling tumor immune interections within micro-environment is quite complex as it consists of heterogenous cellular populations like immune cells, fibroblasts, endothelial cells, extracellular matrix, growth factors, cytokines, healthy cells, and senescence cells. Interaction strengths changes between the sub populations in a healthy and an aging microenvironment. Processes of aging affects tumor initiation and understanding its influence on the tumor microenvironment is challenging [33]. Aging microenvironment also influences tumor progression [34]. Therefore, its important to consider the role of both healthy and aging microenvironment while modeling tumor immune system to understand the dynamics and develop effective therapies. Mathematical oncology has paved the way to develop simple conceptual models (phenomenological, predictive, or mechanistic) and validate the hypothesis experimentally and test predictions on clinical data to improve the efficacy of tumor treatment. Numerous mathematical models have been used to understand these interactions like [35], population dynamics based [36], tumor immune interaction with treatment [37], and anti immune activity by tumor on immunotherapy [38]. Authors also proposed a general framework [39] of modeling the tumor immune interaction [40]. Interaction between cancer cells, immune cells, and healthy tissue cells [41]. Interactions between cancer cells, effector cells, and cytokines (such as IL-2, TGF-*β*, IFN-*γ*) [42], and interactions among cancer cells, effector cells, and naïve effector cells [43]. Four dimensional model consisting of three populations; cancer cell, immune cells and normal cells along with cytokines [44].

A system of three differential equations with possible therapy treatment strategies is a faithful representation to understand the inherent dynamics at the population level. ODE model describing the interaction between the three populations namely tumor (T), immune (I), and senescence (S) is discussed below.

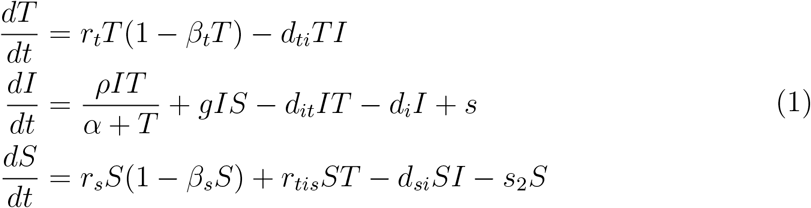

In system of equations 1, T is tumor cells, I is immune cells and S is senescence cells. Consider logistic tumor growth rate *r*_*t*_, competition *β*_*t*_, and tumor clearance rate *d*_*ti*_ by the immune system. Tumor induced immune growth rate *ρ*, tumor saturation term (*α*+T), senescence secreted growth rate *g*, immune death rate by tumor *d*_*it*_, immune apoptosis *d*_*i*_, and immunotherapy term as s. Also consider senescence logistic growth rate *r*_*s*_, senescnce competition *β*_*s*_, therapy induced senescence rate *r*_*tis*_, senescence clearance rate *d*_*si*_ by immune system, and senescence cell death by senolytics as *s*_2_. In the figure 1, three cellular populations exists namely T,I, and S interacting with each other through positive and negative feedback loops.

**Figure 1:**
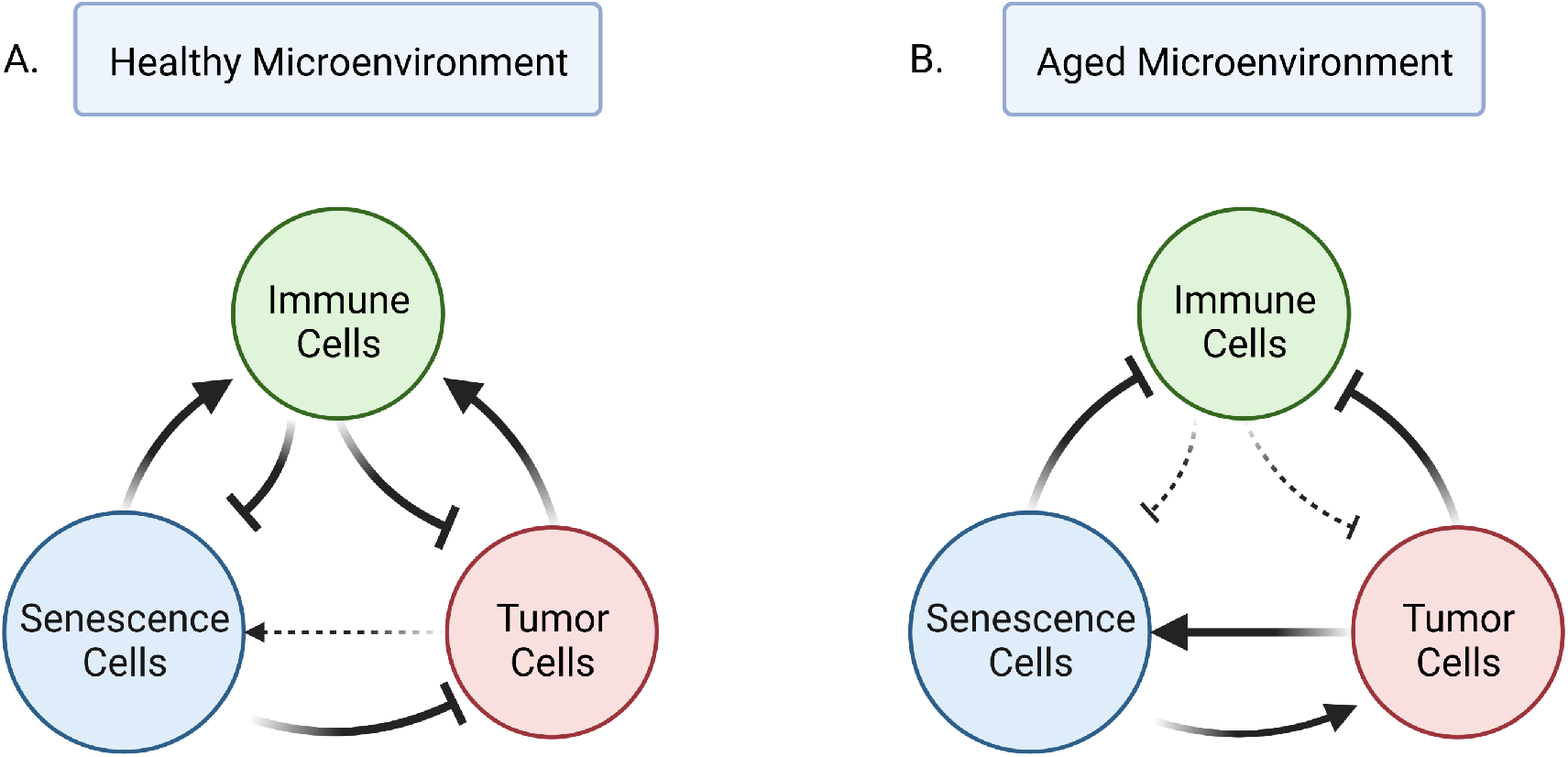
Healthy and Aged Tumor Immune Microenvironment (TME). In a healthy tumor immune microenvironment, senescent cells suppresses tumor growth and recruits immune cells for their clearance. Similarly, tumor cells activate immune cells and immunce cells eliminate tumor cells. In an aged tumor microenvironment, senescence cells modify the immune cells by inhibit their function and promote tumor growth. Here, tumor cells inhibit the immune cells. Senescence cell accumulation beyond a certain threshold may lead to the transition of a healthy to an aged tumor immune microenvironment [6]. Therapy Induced Senescence (TIS) may play a potential role in maintaining a homeostatic tumor immune microenvironment.

Simulation results are present in the figure 3. Fig. 3a shows logistic tumor growth and senescence cell accumulation in the presence of a weak immune system reflecting aged microenvironment. Fig 3b shows tumor decay, explaining strong immune system prevents the growth of tumor and senescence cells accumulation reflecting the clearance in health microenvironment. In 3c, strong senescence signaling (*g*) by secretion of SASP factors loads the immune system results in establishing of tumor population. In 3d, high TIS (*r*_*tis*_) therapy induced senescent tumor cell accumulation reduces tumor burden but may result in tumor relaspe at later stage. Together with senolytics and TIS in 3e shows oscillatory behavior may provide a population level control process by combining therapy strategies as proposed in [14]. And in 3f, TIS activates the innate NK cells with elimination mechanism (high *d*_*si*_) resulting in senescence tumor cell clearance.

**Figure 2:**
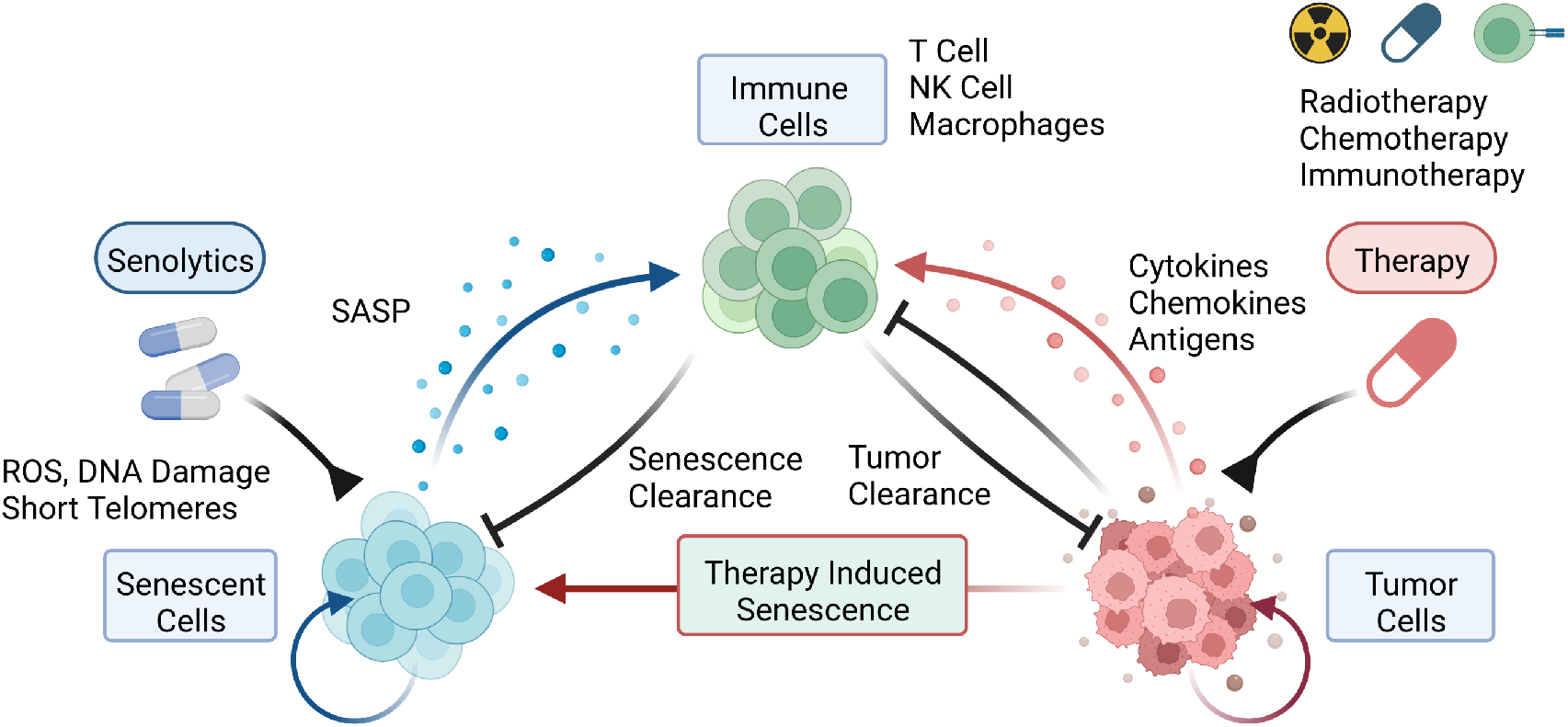
Tumor Immune Senescence Model. Created with BioRender.com. A simplified version of the aging tumor immune microenvironment comprises tumor cells (T). immune cells (I), senescent cells (S), and secreted factors like senescence associated secretary phenotype (SASP), cytokines or growth factors. Senescence cells recruits immune cells and signal them for clearance through SASP. Tumor cells through pro and anti tumor signaling via secretary factors can recruit immune cells for tumor clearance or inhibit immune cells respectively. External agents like senolytic drugs or SASP inhibitors reduce the senescence burden. Therapies such as radiotherapy, chemotheray, or immunotherapy reduces the tumor burden. Therapeutic stress leads to therapy induced senescence (TIS) of tumor cells becoming senescent.

**Figure 3:**
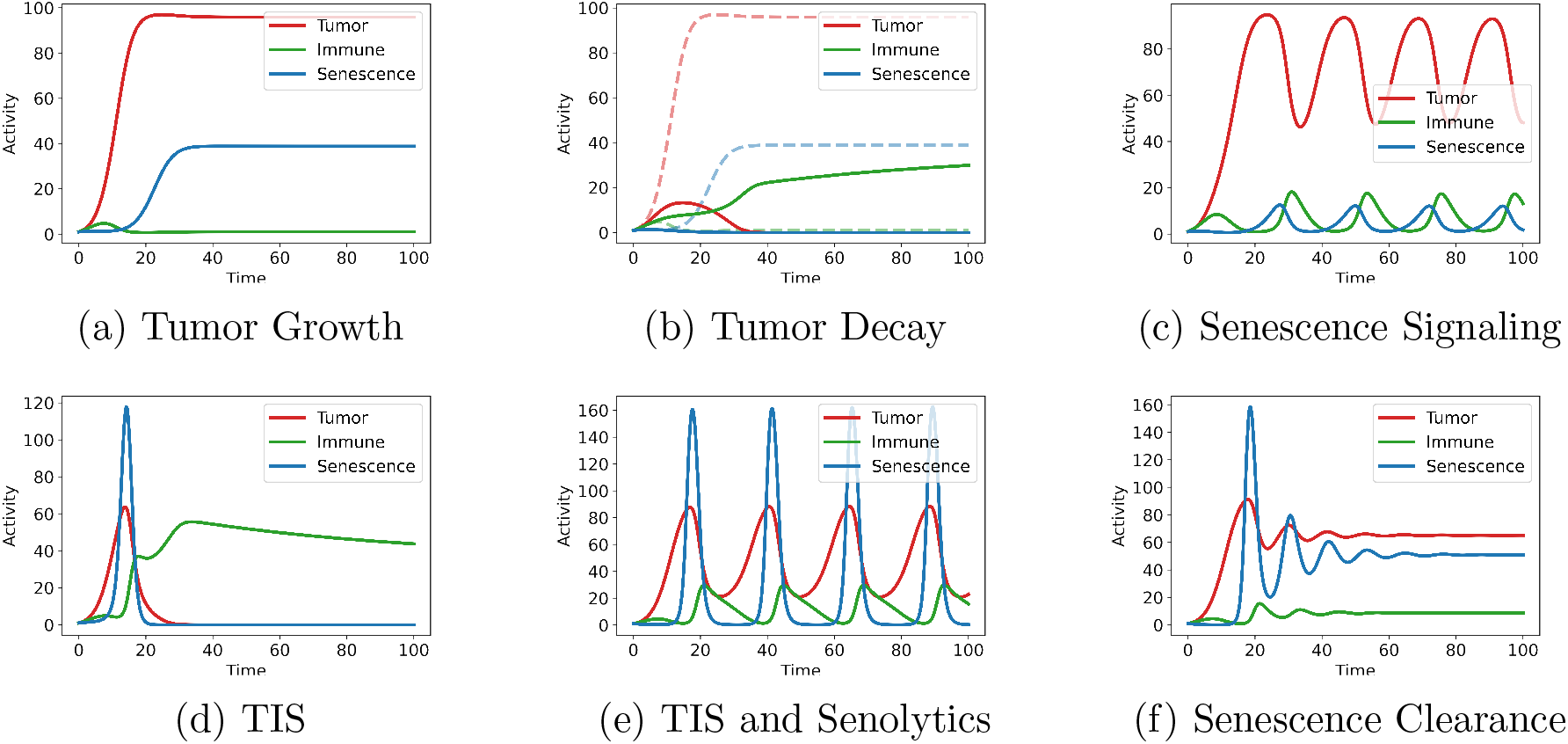
Tumor Immune Senescence Model (A.) Logistic growth of the T and S cell population with a weak immune system in the aged TME. (B.) Strong immune system causes effective clearance which leads to decay of both T and S population in the healthy TME. (C.) S cells recruits I cells causing a loading effect on the immune system which may give rise to a robust build up of T cell population. (D.) TIS can result in effective clearance by the immune system. (E.) Combined TIS along with senolytic drugs may provide a robust population level regulation indicated by a sustained oscillatory behavior. (F.) Senescence tumor cell clearance by the activated NK immune cells shown in a decayed oscillatory manner.

### Stochastic Tumor Immune Senescence Model

TME is highly heterogeneous in nature due to undergoing phenotypic modifications, state transitions and competitive interactions. Tumor comprises sensitive, resistant, and hybrid sub populations. Similarly, macrophages, NK cells, T cells (different types), B cells, etc are present within immune sub population. Phenotypic alterations in early and late stages of senescence cells [30]. Stochasticity changes the therapy temporal dynamics [23]. Hence, adding an inherent noise to the system of equations 1 to closely model the role of TME heterogeneity with and without therapy.

Simulation results are present in the figure 4 which indicate an interesting observation that heterogeneity seen in fig. 4f can maintain robust tumor population despite combination therapies in comparison with the results present in figure 3f.

**Figure 4:**
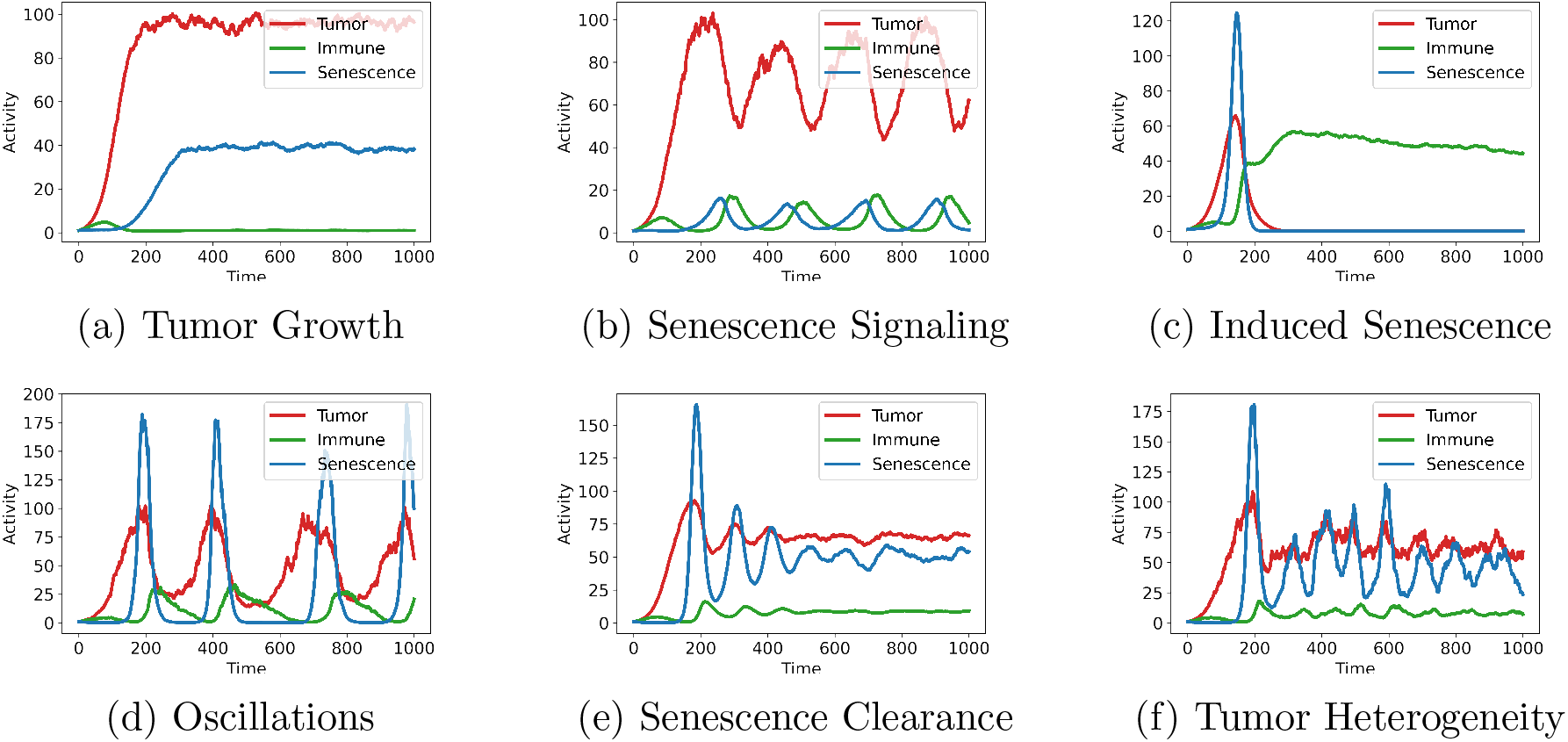
Stochastic Tumor Immune Senescence Model (a) Logistic growth of tumor and senescence cells with weak immune system. (b) Senescence recruiting immune cells. (c) Therapy induced senescence. (d) TIS with senolytics indicating sustained oscillations. (e) Senescence cancer cell clearance by immune cells with decayed oscillatory behavior. (f) High tumor heterogeneity leads to sustained tumor growth despite senescence tumor cell clearance by immune cells.

### Therapy Intensity and Temporal Dosage Regimen

Identifying a correct therapy administration time and dosage intensity is challenging for treating tumors. Here, an attempt is made to model the combined therapy temporal and intensity dynamics and predict low or effective dosage intensities, precise or imprecise dosage timing, and correction of earlier imprecise dosage timing leading to therapeutic shift in the tumor by a later precise dosage timing. For such phenomenological model, parameters are based on trial and error, does not fit true volumetric capacity number of cells in tumor microenvironment and require further validation to fit with an experimental data.

Therapy dosage timing and intensity regimen is present in the figure 5. Figures 5a, 5b, and 5c indicate low, average, and effective dosages respectively. In figures 5d, 5e, and 5f timing interval for precise, imprecise, and correct therapy treatment in eliminating tumor and senescence burden is shown. In figures 5g and 5i, therapeutic shift occurs due to therapy correction for earlier imprecise therapy. Figure 5h indicate ineffective therapy simply reduces the burden without any improvement.

**Figure 5:**
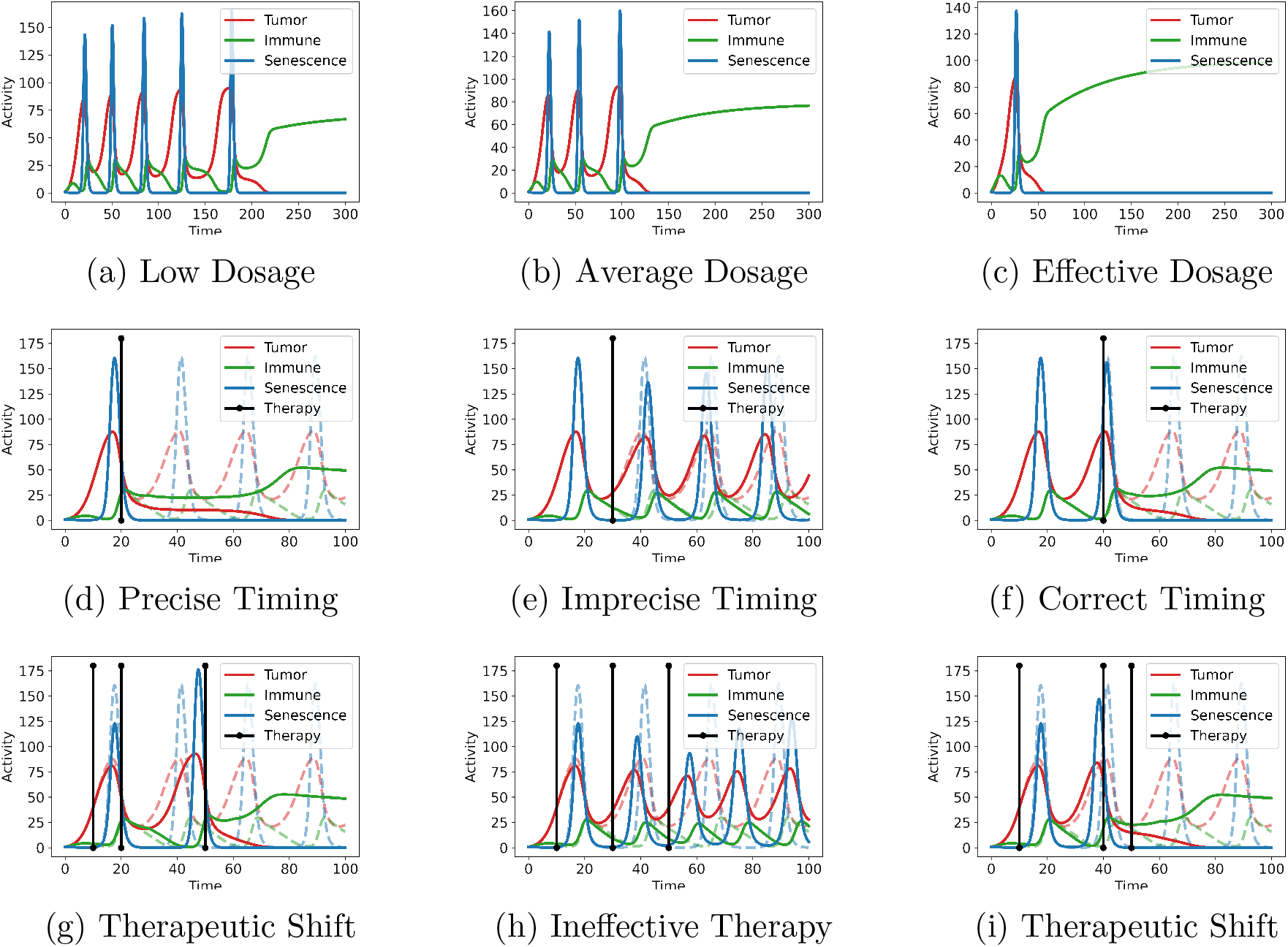
Therapy intensity and temporal dosage regimen. Consider all the simulation results in this figure are in the presence of senolytics. (a) Low therapy dosage takes multiple cycle to eliminate S and T. (b) Normal therapy dosage takes reduced number of cycles for elimination. (c) Effective therapy dosage may require just one cycle for elimination. (d) Precise therapy interval and timing (TI=20) may eliminate T and S in just one shot. (e) Imprecise (TI=30) timing causes temporal shift in population dynamics. (f) Correct therapy interval (TI=40) timing at a later duration can still be effective. (g) Improper therapy at an earlier time interval (TI=10) shifts the therapeutic effect and can be corrected at later stage (TI=50) with additional dosage cycle. (h) Incorrect therapy intervals slightly reduces the burden without improvement. (i) Therapy correction similar to (g) at a later time interval (TI=40).

## Conclusions

Senescent cells within aged tumor microenvironment (TME) modifies the tumor immune interactions pre and post therapeutic interventions. Here, A minimal model tried to explain the mechanisms of senescence tumor cells clearance by the NK cells and senescence cell accumulation threshold hypothesis. Model explains interesting behaviors like tumor and senescence cell growth during weak immune response, tumor decay in strong immune response within healthy TME or by externally induced immune cells, senescence cell signaling supress tumor in healthy TME or promote robost tumor growth in aged TME, therapy induces senescence (TIS) along with senolytics display oscillatory behavior, and senescence cell clearance by natural killer (NK) cells shows oscillatory decay behavior. Inherent heterogeneity and noise can maintain tumor despite therapies. Lastly, model provides preliminary analysis for therapy dosage intensity and timing regimen leading to therapeutic shift of tumor.

Model can be used for the development of patient-specific personalized tumor treatment plans using mathematical modeling of tumor immune interaction by understanding the therapeutic efficacy differences within healthy and aged tumor microenvironment. Might be a key to potential benefits [27] despite developing cell-based therapies. A conceptual model involving patient’s treatment response to develop clinical dosage plans using therapy intensity and temporal regimes in the clinic can improve the efficacy of immunotherapies treatment by transitioning from therapeutic tool to therapeutic effect [45]. For example, combination of Dasatinib and Quercetin reduced senescent cell burden in mice and demonstrate the efficacy of senolytics for reduced frailty [46]. Future modeling approaches must consider the role of aging in modifying the tumor microenvironment for possible such two punch therapies [14] or adaptive therapies [19] to be effective in regulating tumor. Similar to immunoediting phases of immune cells, there could be a senoediting kind of features for senescence induction, maintenance, and escape [11] which can help in discovering new ways to treat cancers combining immunotherapies and senotherapies.

## Acknowledgement

This work is a part of the iCurious.in organisation to promote curiosity driven research. I personally thanks all the referenced researchers for providing the foundation for conceptualization, formalization, modeling, simulation, and writing this manuscript.

